# Homomorphic Encryption: An Application to Polygenic Risk Scores

**DOI:** 10.1101/2024.05.26.595961

**Authors:** Elizabeth Knight, Jiaqi Li, Matthew Jensen, Israel Yolou, Can Kockan, Mark Gerstein

## Abstract

1

**Background:** Polygenic risk scores (PRSs) have emerged as a powerful tool in precision medicine, enabling personalized risk assessments for complex diseases. However, the use of sensitive genomic data in PRS calculations raises concerns about privacy and security. Fully homomorphic encryption (FHE) offers a promising solution by allowing computations on encrypted data, preserving the privacy of both genomic information and PRS models.

**Methods:** Here, we present an application of FHE for encrypted PRS calculations using a particular protocol (CKKS) within the Lattigo library. Our approach involves a three-party system: clients (clinicians handling sensitive genetic data), modelers developing a PRS (academics or companies), and evaluators (a local hospital running the models while maintaining data confidentiality). We demonstrate the feasibility and accuracy of our approach by applying it to synthetic datasets of various sizes and to a robust 110k–single-nucleotide polymorphism (SNP) model for schizophrenia. The complete codebase and a sample dataset are available at https://github.com/gersteinlab/HEPRS.

**Results:** The difference between traditional plaintext and encrypted PRS calculation results is negligible: the *R*^2^ is 0.999 and the mean squared error is 2.27 *×* 10^−6^. Moreover, while the encrypted calculation is roughly 1,000 times slower than conventional non-encrypted ones (when considering only the core PRS calculation), the computation remains feasible on a single-CPU node. For example, processing ∼1,100 individuals with ∼110k SNPs took six minutes and ∼65 GB of memory on a laptop computer. In addition, we investigated the impact of the encryption parameters on the computational time and accuracy in detail, showing the expected slowdown with higher security settings.

**Conclusion:** Our approach showcases the applicability and feasibility of using FHE on real-world PRS models. With the pressing need for privacy-preserving solutions in the era of precision medicine, our work serves as a pilot application, offering a simple use case and providing a detailed comparison and evaluation in terms of accuracy, cost, and scalability.

## 2 Introduction

Advances in genetic sequencing technologies have significantly reduced both time and cost, making sequencing a commonly used practice in biomedical research and precision medicine [1]. Genomic sequencing data have the potential to provide valuable insights into biological processes and clinical treatment, especially with the multitude of associations established between genomic loci and traits or diseases through genome-wide association studies (GWAS) [2]. Such studies hold promise for various clinical applications, from more accurate molecular diagnosis to drug target prioritization [3] [4]. A meaningful clinical advance stemming from GWAS is the development of polygenic risk scores (PRSs), which are statistical models that utilize genetic variants to predict an individual’s risk of having a particular trait or condition, such as disease, height, or intelligence [5]. PRSs have the potential to bring personalized approaches to health and wellness, from early detection and prevention through lifestyle changes or medical interventions to more effective and individualized care. PRSs can also identify subgroups of individuals more likely to respond to certain medications or treatments, improving patient outcomes [6]. For example, a recent study found that PRSs can indicate future heart attack risk, prioritizing patients for preventive statin treatments [7]. Similarly, PRSs for breast cancer can identify subgroups of individuals with high predicted incidence for early screening interventions [8], further highlighting the utility of PRSs in a clinical setting.

GWAS for most common traits (such as height) and diseases (such as breast cancer) typically require genomic data from tens of thousands of individuals at hundreds of thousands of loci for accurate results [9] [10]. The sheer amount and complexity of these genetic data can strain local computational resources and often require the involvement of third parties or cloud-based infrastructures, thereby increasing the risk of genetic privacy violations [11]. For example, the UK Biobank contains more than 400k samples, whose data are often too large to be fully downloaded, requiring the use of the DNANexus platform for downstream analysis [12]. Protection of patient genomic data is particularly important for PRS calculation, where an individual’s genome is compared against summary results derived from a large, often external cohort. In the face of quantum computing advancements, quantum-secure encryption has become important in ensuring the privacy of data transfers across systems [13]. As genomic data become increasingly important in a healthcare setting, hospitals must adopt advanced encryption solutions to ensure the privacy and security of patient data, especially genomic data, while also reducing costs and improving the efficiency of clinical research. Thus, advanced encryption methods are needed to protect sensitive genomic data from unauthorized access and misuse while allowing physicians to calculate and access their patients’ PRSs securely and confidentially. With the increasing use of PRS models trained on confidential patient information, establishing model security is also critical to protect clinical data and prevent potential model inversion attacks [14].

Despite the potential of PRSs in clinical applications, ensuring the privacy of genomic data remains a challenge. Various methods have been proposed to address genomic privacy concerns, including cryptographic techniques such as homomorphic encryption. Previous applications of fully homomorphic encryption (FHE) in healthcare have set the stage for advancing patient data privacy. For example, [15] initially leveraged this technology to securely handle vital sign data from medical devices. Later, [16] extended the use of FHE to protect genetic data within hospital systems, enabling researchers to access GWAS summary statistics without compromising patient privacy. Naveed et al. [17] provided a comprehensive survey of privacy-preserving methods in genomics, highlighting cryptographic approaches to enhance data security. Similarly, Bonomi et al. [18] discussed the privacy challenges in genomic data sharing and suggested homomorphic encryption as a promising solution for secure computations on genomic data, although they did not implement it in their study. Previous work has also applied homomorphic encryption to genomic data analysis tasks other than PRS calculation. Kim and Lauter [19] utilized the Brakerski-Gentry-Vaikuntanathan (BGV) and Yet Another Somewhat Homomorphic Encryption (YASHE) homomorphic encryption schemes to securely compute minor allele frequencies and chi-squared statistics in GWAS, as well as to calculate Hamming and approximate edit distances between DNA sequences. Their experiments with datasets containing up to 5,000 DNA sequences demonstrated the feasibility of using homomorphic encryption in genomic analysis, although PRS calculations were not performed. In clinical settings, McLaren et al. [20] used homomorphic encryption to securely evaluate specific genetic loci associated with drug resistance or treatment response in HIV patients. This approach pioneered privacy-preserving genomic testing but did not extend to PRSs. Blatt et al. [21] built a privacy-preserving pipeline for large-scale GWAS using an optimized variant of the CKKS homomorphic encryption scheme. The scheme allows researchers to obtain GWAS summary statistics without ever seeing an individual’s genomic data, as the encrypted genomic data reside only in the cloud. The authors noted that PRSs can be computed once the odds ratios are decrypted, but a fully end-to-end encrypted pipeline for PRS was left for future work. Implementing FHE in PRS calculation could significantly enhance data security by maintaining patient genomes in encrypted formats, potentially reducing vulnerabilities and costs when facilitating the secure transfer and manipulation of encrypted genomic data.

Homomorphic encryption is a promising solution to the problem of preserving confidentiality while enabling important computations on sensitive genomic data. This type of encryption allows third parties to perform computational functions on encrypted data (ciphertext) without first decrypting it. It addresses privacy concerns in digital communication by preserving confidentiality during data manipulation without exposing it to unintended parties [22]. There are several types of homomorphic encryption, including partially homomorphic, somewhat homomorphic, leveled fully homomorphic, and FHE, each offering varying degrees of computation on encrypted data [22]. Many commonly used homomorphic encryption schemes rely on the Ring Learning With Errors (RLWE) problem, which deliberately introduces a small, controlled amount of noise into ciphertexts to guarantee security as each computational operation is performed (with addition contributing less noise than multiplication). The noise accumulates, and the RLWE framework ensures that this noise cannot be reversed or “subtracted out” without the secret key, thereby preventing backward reconstruction of the original plaintext. However, different types of homomorphic encryption have limitations regarding the set of circuits, or the range and complexity of computational sequences that can be performed on encrypted data, and the types of gates, or basic operations (such as addition or multiplication), that are supported in these computations [23] [24].

Here, we apply FHE specifically to genotype data for secure PRS calculation and phenotype prediction. Utilizing the CKKS protocol for FHE within the Lattigo library [25], we present a novel approach that allows for the computation of PRSs directly on encrypted genetic datasets, obtaining results securely and privately. Our approach entails the propagation of encrypted data across three parties (clients with sensitive genetic data, modelers with existing PRS models, and evaluators to interpret PRS findings), preserving both model and genomic privacy when communicating PRS results back to patients [26]. While other studies have proposed homomorphic encryption of PRS models and provided proof-of-concept on limited artificial datasets [27], we take advantage of the scalability of FHE to preserve the privacy of robust PRS models that contain a larger number of significant single-nucleotide polymorphisms (SNPs). In addition to robust assessment with synthetic datasets to show how our model scales with increased SNP numbers, we apply our FHE-based protocol to a 110k–SNP PRS model for schizophrenia [28], which demonstrates accuracy in predicting schizophrenia risk in a cohort of over 1,200 individuals, with minimal decrease in performance compared to non-encrypted PRS. We also thoroughly investigate the trade-offs associated with encryption parameters that influence computational accuracy, memory, and time for large PRS models, which are relevant for real-world healthcare applications. Our FHE protocol can preserve PRS while operating within a reasonable timeframe; in fact, evaluating the PRS of one individual with the 110k–SNP model required only 4.9 seconds and 3.3 GB of memory with optimal parameters, while calculations for 1,000 genotypes took less than five minutes and used 130 GB of memory on a conventional laptop (Macbook Pro 2021). By using FHE to obtain meaningful results from realistic PRS models while maintaining the security of the underlying genomic datasets, our study provides valuable insights into the potential applications of FHE in genomics and healthcare. An implementation of our FHE protocol is available as an open-source software package called HEPRS at https://github.com/gersteinlab/HEPRS.

## 3 Methods

### 3.1 Method Overview

Our work establishes a novel privacy-preserving protocol utilizing homomorphic encryption to predict phenotype risk from genomic data by involving three independent parties: the client, the modeler, and the evaluator (Fig. 1). Our protocol delineates distinct roles—clinicians could be clients providing genomic data, researchers or private companies could serve as modelers creating the PRS, and hospitals or centralized systems could act as evaluators processing the encrypted genomes. This approach ensures effective use of genomic information while prioritizing patient confidentiality.

**Figure 1:**
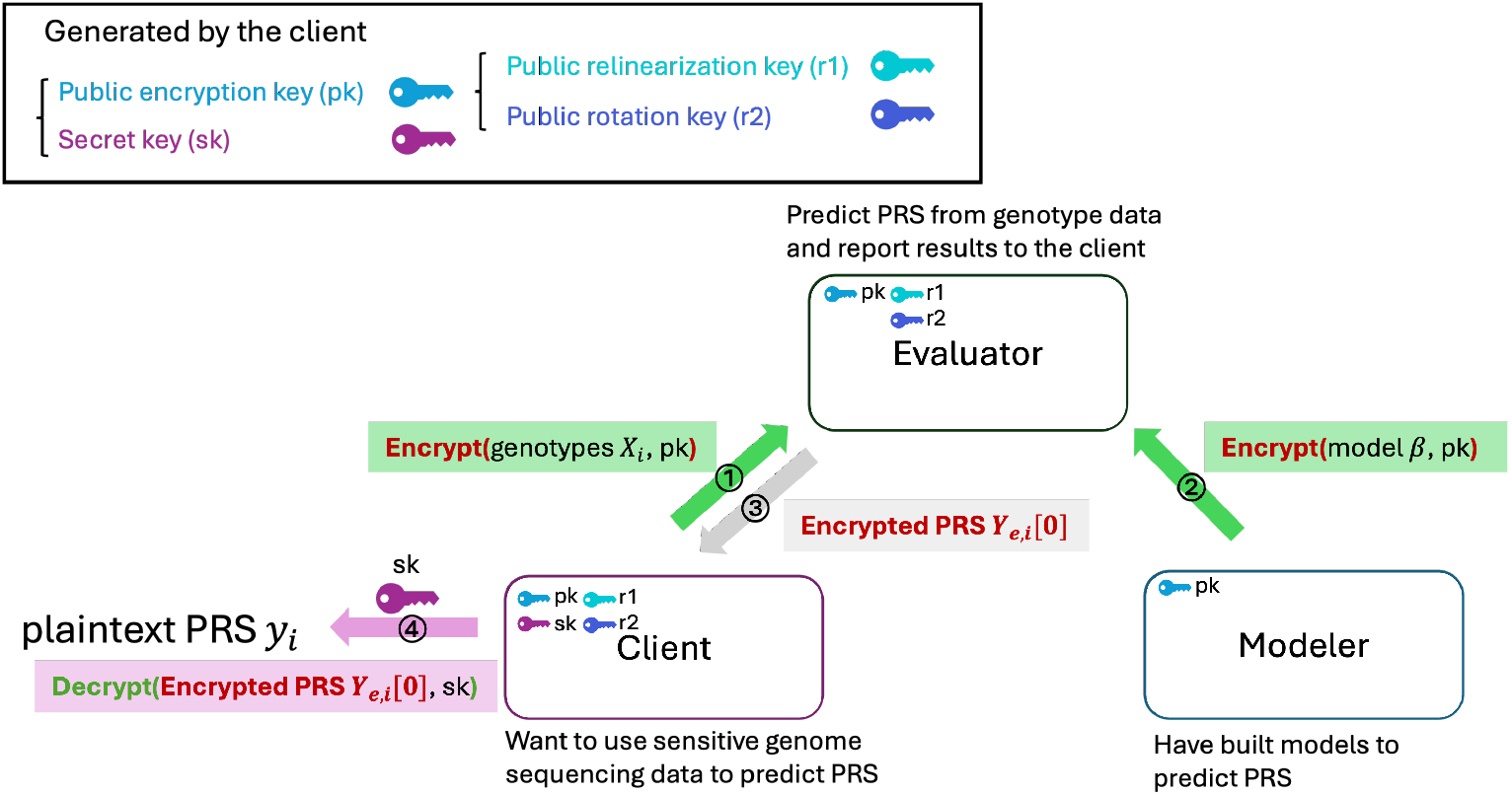
Schematic illustration of the encryption setting. The public encryption key (pk), public relinearization key (r1), public rotation key (r2), and secret key (sk) are generated by the client. The public keys (pk, r1, r2) are shared with all three parties, while the secret key is only known to the client. The client performs homomorphic encryption of their sensitive genetic data using the public key, and the modeler encrypts their model using the same public key. The clients’ genomic data and the model are sent to the evaluator in an encrypted form. The evaluator cannot decrypt the data or the model without the secret key. However, the evaluator is able to evaluate the genetic data and obtain model predictions, which remain encrypted. The encrypted phenotype predictions are reported back to the client, who can decrypt the data using the secret key and read the predictions.

This protocol utilizes the CKKS encryption scheme [29], which is quantum-secure and allows for a broader range of bootstrapped computations on encrypted data than other homomorphic encryption methods. Using both synthetic and real genomic datasets, we showed that our encryption scheme efficiently calculates PRS values without sacrificing privacy or accuracy compared with traditional non-encrypted PRS methods. We further systematically analyzed the various encryption parameters used in our method to report their effects on accuracy (R-squared and area under the receiver operating characteristic curve [AUROC]), memory usage, and computational time. Overall, our findings underscore the practicality of employing FHE in clinical genomics, particularly for PRS calculations where data privacy is paramount, without compromising computational integrity.

#### 3.1.1 Evaluation Strategy

The FHE encryption framework involves the client, the modeler, and the evaluator [26] (Fig. 1). The roles within our protocol can be adapted to various configurations, aligning with the operational structures of clinical genomics environments. For instance, the client could be a clinician tasked with diagnosing or treating a patient, ensuring that the genomic data are utilized effectively while maintaining patient confidentiality. The modeler could represent a centralized data repository, akin to the UKBiobank resource, which manages PRS calculations and the underlying summary statistics for various disorders. The evaluator could be a hospital’s biobank, which can securely access and process a patient’s encrypted genome in conjunction with their electronic health record. In this case, the client has genomic data and wants phenotype predictions, the modeler trains a model for predictions, and the evaluator evaluates the model using the client’s data and reports the forecast to the client.

Privacy is protected throughout the framework using homomorphic encryption, where the client’s data and model are encrypted, and computations (i.e., calculating the PRS) are performed only in an encrypted form (Fig. 1). Specifically, the client first generates a public key and a secret key and encrypts the data using the public key. The client then shares the public key with the modeler and the evaluator, and the modeler encrypts their model using the same public key. The client and modeler both share their encrypted data with the evaluator. The evaluator performs the computation using the encrypted model, encrypted data, and public key. Finally, the evaluator returns the encrypted predictions to the client, who then decrypts the results with their secret key. We assume no cross-talk between the client and the evaluator. It is also pertinent to acknowledge that the modeler, in handling GWAS summary statistics, is a custodian of sensitive information, as minor allele frequencies could allow for identifying individuals in the study [30]. In this context, the modeler must trust that the evaluator will not share the encrypted model with the client. In this way, all data are encrypted and only the client can access the plaintext predictions (Fig. 1).

### 3.2 Technical Overview of Key Aspects of FHE for PRS

We use the CKKS encryption scheme, implemented in the Lattigo library [25], as the FHE method in our encrypted PRS calculation framework. The CKKS scheme gains NP-hard security by leveraging the RLWE problem. Homomorphic evaluation allows the scheme to perform a specific set of operations on ciphertexts, such as addition and multiplication, and produce a new ciphertext as a result. The CKKS scheme is classified as lattice-based cryptography and is quantum-secure [29]. Our parameter set meets 2^128^ classical and quantum gate complexity, satisfying NIST category one security[31]. Although lattice problems do admit a quadratic speed-up in quantum models, this does not translate into a large reduction in security levels. As shown in the Homomorphic Encryption Standard [32] and the CIC/IACR guidelines [33], the difference between classical and quantum estimates is only a few bits. For example, parameters yielding ∼128 bits of classical security typically yield 124-127 bits quantum, not a large drop.

Table 1 outlines each important parameter and its meaning in the context of the encrypted PRS pipeline. We first encode the model *β* and the genotype vector of each individual *X*_*i*_ for *J* individuals as CKKS plaintexts; CKKS packs a real vector *X*_*i*_ into a polynomial of degree N with coefficients in the ring 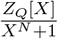 (this is also stored in vector format *X*_*e,i*_). These plaintexts are then encrypted to produce the ciphertext matrices used in the subsequent homomorphic computations (see Methods 3.6 for details).

**Table 1:** N is the ring dimension, which determines the slot capacity. The modulus budget QP allows for more depth of computation and is bounded by security for a given N. Slots represents the maximum of N/2 values that can be encoded at once, or in other words, the maximum vector size for the given ring dimension. M is the number of SNPs included in the PRS calculation. J is the number of individuals tested. K is the number of encrypted vectors that comprise one encrypted genome 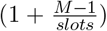. X_*i*_ is the genomic vector for individual i, and ***X***_***e***,***i***_ is the encrypted genomic matrix for individual i. β is the vector containing model weights for the PRS and ***β***_*e*_ is the model matrix with encrypted weights. Y_*e,i*_ is the encrypted results vector and y_*i*_ is the unencrypted result scalar that communicates an individual’s risk for disease.

| Data type | Symbol | Description | Dimension |
| --- | --- | --- | --- |
| Input data | $M$ | number of SNPs included in PRS model | scalar |
| | $J$ | number of individuals | scalar |
| | $X_i$ | genomic data vector | $(M)$ |
| | $\beta$ | unencrypted model vector | $(M)$ |
| | $security$ | chosen security NIST category | |
| Dependent on security + $M$ | $QP$ | extended modulus (allows for more depth of computation with security compromise) | scalar |
| | $N$ | ring dimension (determines slot capacity and chosen to meet target security for given $q_{max}$) | scalar |
| | $slots$ | maximum vector size $(N/2)$ | scalar |
| | $K$ | number of encryption vectors per one genome $M/(N/2)$ | scalar |
| Calculated values | $X_{e,i}$ | encrypted genomic matrix | $(K, N/2)$ |
| | $\beta_e$ | encrypted PRS model matrix | $(K, N/2)$ |
| | $Y_{e,i}$ | encrypted result vector | $(N/2)$ |
| | $y_i$ | unencrypted result | scalar |

Figure 2 outlines how we encode the genomic vector into the encrypted domain to prepare for FHE evaluation. During encoding, we break each individual’s genome vector *X*_*i*_ with dimension 1*×M* into a matrix of dimension *K×N/*2. This becomes matrix ***X***_***e***,***i***_ with dimension *K×N/*2. After performing this process for each individual, we generate a three-dimensional tensor of dimension *J × K × N/*2. We repeat this process for vector *β* to obtain matrix ***β***_***e***_ with dimension *K* x *N/*2. We ultimately compute the calculation *X*_*i*_ *× β* = *y*_*i*_, where *y*_*i*_ is the scalar output from the PRS model. We outline our algorithm in Algorithm 1 with a greater explanation of the individual steps in Methods Section 3.6.

**Figure 2:**
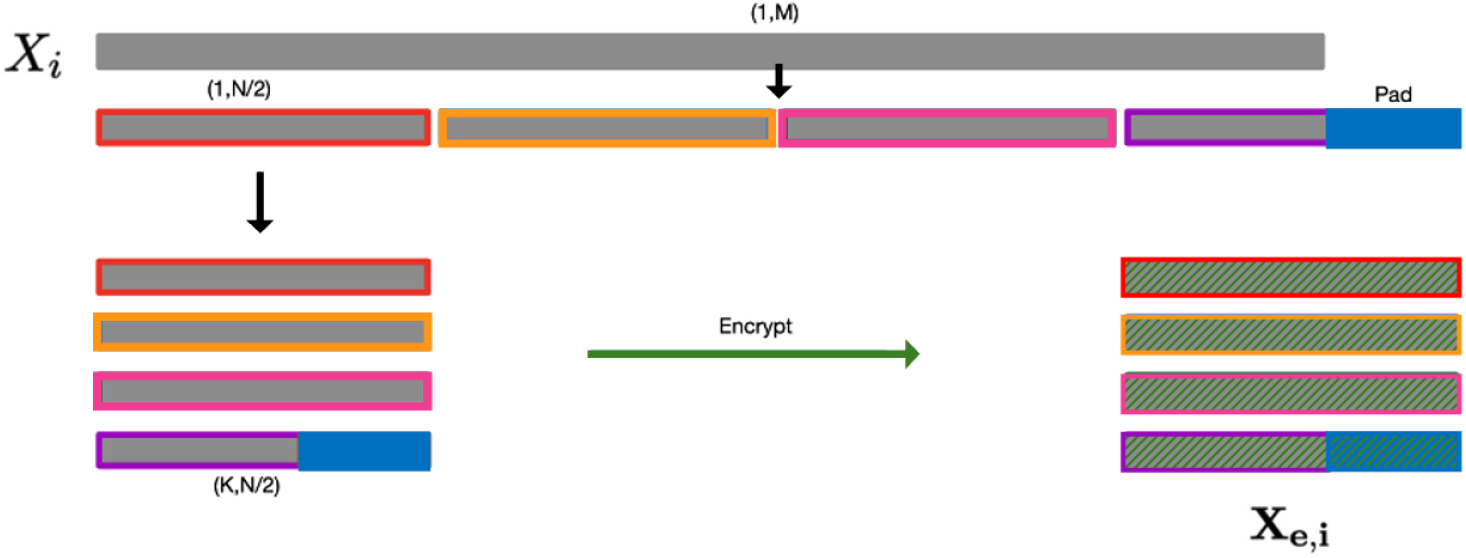
This schematic shows how we encode the genomic vector X_*i*_ into a genomic matrix in the encrypted domain X_*e,i*_. Green hash denotes encrypted. Blue denotes the padding.

In Table 2, we highlight the important parameters that customize the CKKS encryption scheme

**Table 2:**
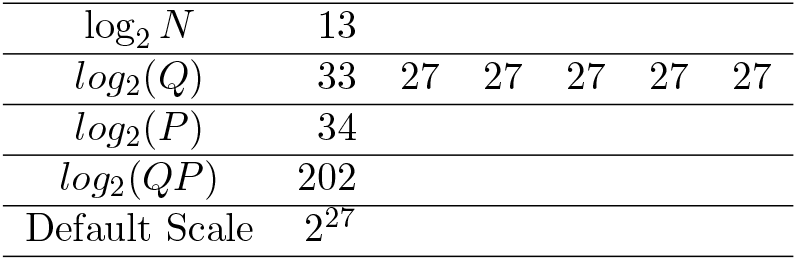
Explanation of encryption parameters. We work in the cyclotomic ring R = ℤ_*Q*_[X]/(X^*N*^ + 1). Here N is the ring dimension; Q = ∏_*i*_ q_*i*_ is the ciphertext modulus (each q_*i*_ ≡ 1 mod 2N); and P is an auxiliary prime used during key switching and relinearization. In our setup, log_2_ N = 13 (N = 8192), log_2_(q_*i*_) ∈ {33, 27, 27, 27, 27, 27 }, and log_2_(P) = 34, so the largest modulus used at any point is q_max_ = QP = 2^202^. Security for RLWE based HE is assessed against N and the largest modulus q that appears during the computation. With N = 8192 the HE Standard’s 128bit recommendations allow roughly log_2_(q) = 218 for classical and log_2_(q) = 202 quantum [31]. The default scale specifies the amount to scale the plain-text during the encoding process, which impacts the precision and maximum depth. We use a pretested parameter configuration specified by the Lattigo package.

#### Algorithm 1

Multiply vectors and add together

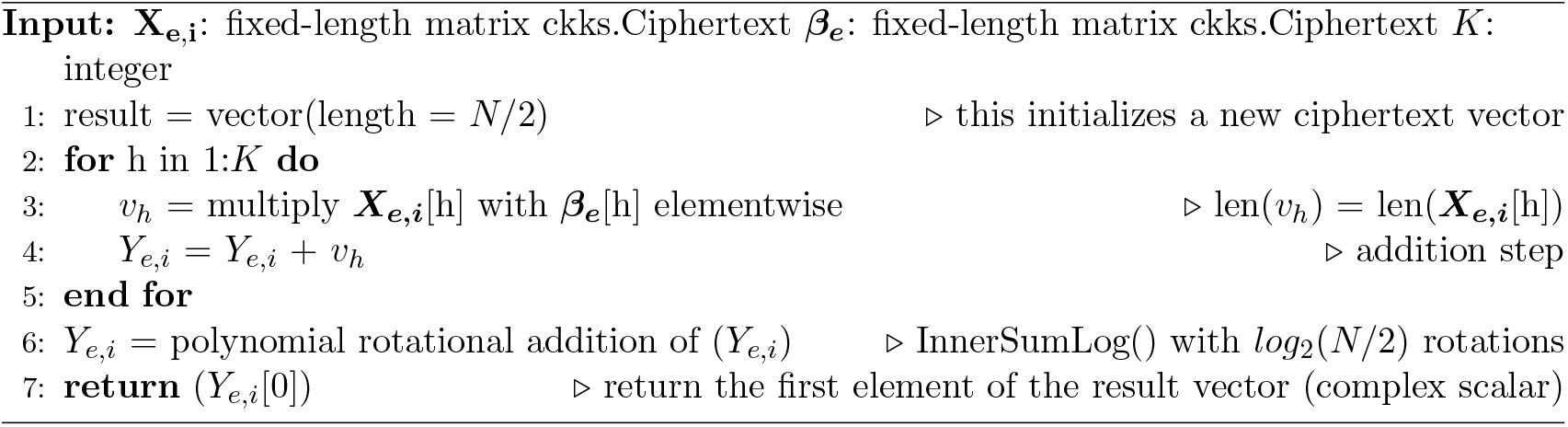

for optimal results for a given task. The parameter *N*, in the context of CKKS (and other homomorphic encryption schemes), represents the degree of the polynomial ring over which the encryption operates. CKKS encodes vectors into polynomials of a degree defined by *N* (we provide further explanations in the Methods section). The ring dimension *N* is chosen to satisfy a greater than 128-bit classical security for a given *log*_2_(*Q*). The value of *N* is typically chosen to be a power of 2 for efficient implementation, mainly because fast Fourier transform algorithms, which are used for polynomial multiplication, are most efficient when operating on sequence lengths that have a power of 2. The values of *Q* and *P* determine the lowest possible value of *N* to maintain security (as larger N increases computational cost). The genomic vectors are converted into a matrix when encoded into the encrypted domain defined by a cyclotomic polynomial, whose domain is defined by *Q* and *N. Q* determines the precision of encrypted genetic data, ensuring accurate PRS computation without loss of data integrity. *P* is critical during the relinearization step post-multiplication to safeguard the computation’s accuracy. We describe each parameter in Table 2.

### 3.3 Key Generation

The key generation step is implemented using the Encrypt input function in the Lattigo library. In this function, we employ the CKKS parameters to create a new key generator through the “NewKeyGenerator” method provided in the CKKS library within Lattigo. The “GenKeyPair” method generates a public-private key pair (pk and sk), and the “GenRelinearizationKey” method creates the relinearization key that enables efficient computation on encrypted data. This method requires the secret key and max level parameters, the latter of which we set to 2 (to support addition and subtraction calculations). Finally, the “GenRotationKeysForInnerSum” method generates rotation keys, which allow operations to include rotations of encrypted data (for instance, during the inner product calculation). The function returns the generated keys and parameters for further use in the encryption and evaluation steps of the homomorphic encryption scheme.

### 3.4 Implementation of Homomorphic Encryption

To implement FHE, we first load and encrypt the genotype data on the client side. During this step, the client generates the secret decryption key (*s*_*k*_), which is then used to derive the public encryption key (*p*_*k*_), the public relinearization key (*r*_1_), and the public rotation key (*r*_2_). These public keys are shared with all parties. The client then encrypts the genotype data (*X*_*i*_) to transform each individual’s genotype into a two-dimensional ciphertext matrix (*X*_*e,i*_). The chosen parameters determine the dimensions of the vector.

Next, the modeler loads and encrypts the PRS model *β* using the public encryption key (*p*_*k*_) provided by the client. A similar scheme, dependent on the parameters, saves each model as a two-dimensional ciphertext matrix *β*_*e*_.

After this, the evaluator loads and evaluates the encrypted model *β*_*e*_ on the encrypted genomes *X*_*e,i*_. In this arrangement, we assume that the evaluator and client are both trustworthy and cannot collude; the encrypted model is never shared with the client. The evaluator uses the public relinearization key *r*_1_ and rotation key *r*_2_ provided by the client, along with the encrypted model *β*_*e*_ and genome *X*_*e,i*_, to perform a polynomial evaluation and obtain the encrypted phenotype (i.e., PRS value *Y*_*e,i*_[0]). This step involves an inner product calculation performed using homomorphic encryption, as detailed below.

Finally, once the client receives the encrypted phenotype result *Y*_*e,i*_[0] (risk score for disease), the client decrypts the result with the secret key *s*_*k*_, producing the final PRS *y*_*i*_.

We implemented the protocol using Golang. Golang’s suitability for cryptography is underscored by its comprehensive standard library, which includes support for common cryptographic algorithms, along with features such as concurrency support, inherent memory safety, and strong typing with compile-time checks.

### 3.5 Implementation of Inner Product Using Homomorphic Encryption

We first encode both the model *β* and *X*_*i*_ genome for *J* individuals using the CKKS plaintext encoding scheme. During encoding, the integer-based plaintext vector *X*_*i*_ (representing genotype values of 0, 1, or 2) is encoded into the domain 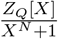. This requires us to represent our vectors as polynomials where all coefficients are ∈ *Z*_*Q*_ and all operations are performed as mod *X*^*N*^ + 1.

In the following section, we focus on the calculation of a single individual’s PRS. Here, the encrypted genomic data matrix ***X***_***e***,***i***_ encapsulates the genotypes, while the model matrix ***β***_***e***_ holds the beta values. Both matrices have dimensions *K × N/*2, aligning with the encryption parameters outlined in Figure 2.

Algorithm 1 is instrumental in the PRS calculation by implementing the dot product of encrypted vectors. Within the for-loop, each row vector from the genomic data matrix ***X***_***e***,***i***_ is element-wise multiplied with the corresponding row from the model matrix *β*_*e*_ using the MulRelinNew() function from Lattigo[25]. This multiplication yields new vectors, *v*_*h*_ = ***X***_***e***,***i***_[h]\****β***_***e***_[h], representing the weighted contribution of each genotype to the PRS (here * represents an element-wise multiplication). The resulting vector aggregates these products, accumulating the combined effect of genotypes and beta values on the PRS. The Lattigo InnerSumLog() function subsequently sums the polynomial terms within the result vector, reflecting the final summation of individual genetic risk factors. The InnerSumLog() operation is called once at the end, outside of the loop with *log*_2_(*N/*2) rotations. This process effectively translates the encrypted genomic information and model predictions into a single PRS value for each individual. This function fails to perform some specialized processes currently executed by more advanced calculation methods such as the plink2 clump function [34]; however, this process calculates a sufficient PRS that differs from the Plink 2 model by only 0.000129%.

### 3.6 Synthetic Data Generation and Ridge Regression

Hapgen2 [35] is a simulation method that resamples known haplotypes quickly and efficiently to produce samples with linkage disequilibrium (LD) patterns that mimic those in real data. Hapgen2 is based on the Li and Stephens model [36] of LD. The method takes in a reference panel of haplotypes as input and generates genotype encodings based on the haplotypes. We used Hapgen2 to simulate genotype encodings for 40,000 individuals across a varying number of SNP sizes, (30k, 50k, 100k, and 130k), based on the 1000 Genomes project reference panel. We later split these individuals into training and testing groups for ridge regression of sizes 30,000 and 2,000, respectively, for each SNP size.

We considered a high-dimensional regression framework for polygenic modeling and prediction

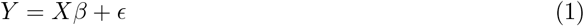

where Y is an *N ×* 1 vector of phenotype values with *Y*_*i*_ ∈ ℝ and N is the number of individuals in the dataset. The parameter *X*^*N*×*M*^ is the genotype encoding matrix with *X*_*im*_ ∈ {0, 1, 2} and M is the number of SNPs in the genotype encoding. The corresponding effect size of each SNP *m* is *β*_*m*_. The parameter *ϵ* is a vector of residual effects for each individual. We are interested in estimating *β*. We assign a Gaussian prior on each *β*_*m*_.

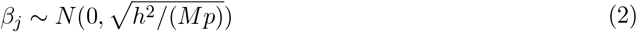

where *p* is the percentage of SNPs with non-null effect size, and *h*^2^ is the total SNP-heritability. Hence, we assume that each SNP with a non-zero effect size explains an equal portion of the total SNP-heritability. To model the impact of SNPs with zero effect size, we select *M*(1 − *p*) random SNP effect size locations in the *β* vector and set the value at those indexes equal to zero. Each of the three artificial phenotypes were generated under the assumption that approximately 90% of the SNPs had non-zero effect size, or *p* = 0.9, a level of sparsity generally consistent with realistic human data [37]. The three levels of SNP heritability used were *h*^2^ = 0.3, 0.6, 0.9.

We used multiple SNP sizes, (30k, 50k, 100k, and 130k), for our simulation data to show that the FHE scheme would apply well to real-world settings where the SNP size varies depending on which traits are being studied. Both values of *p* and *h*^2^ were fixed across the training and testing populations to maintain consistency across the two data populations. The population count of the training dataset was *N* = 30, 000, and for the testing data the count was *N* = 2, 000.

Recall that the estimator of Ridge Regression 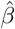 is equal to the mean of the posterior distribution under a Gaussian prior. Hence, the choice of a Gaussian prior on *β* in this PRS setting naturally leads us to use ridge regression. Moreover, an extra column of 1s was added for the intercept term. We note that ridge regression is one of many models researchers use for calculating PRS. More advanced models, such as Bayesian hierarchical models using Gibbs sampling or variational inference, may be more accurate because they have relaxed assumptions on the parameters of the training data. Although we sacrifice computational accuracy in using ridge regression, we gain faster time complexity. In real-world settings, this trade-off is not as important since the model is built once off of training data and not repeatedly calculated. It is important to note that the sacrifice on accuracy is made in the model selection and not in the FHE scheme. Hence, the FHE scheme’s performance should be judged purely on the accuracy of the decrypted prediction values compared to the ciphertext prediction values.

### 3.7 Schizophrenia PRS Generation

We calculated schizophrenia PRSs for 1,146 individuals from the PsychENCODE Consortium study, including 493 individuals diagnosed with schizophrenia and 653 controls [38]. Imputed genotypes and metadata for these individuals were downloaded from the PsychENCODE Consortium data portal [38], with more than 4.3M imputed SNPs available for each individual. Schizophrenia GWAS summary statistics were selected from a recent Psychiatric Genomics Consortium (PGC) study of more than 320,000 individuals and were downloaded from the PGC portal [28]. Using these inputs, we implemented a standardized data processing pipeline for calculating PRSs based on best practice guidelines [39]. We first performed quality control filtering on the GWAS summary statistics by removing SNPs with INFO scores less than 0.8, as well as ambiguous and duplicate SNPs. Next, after lifting over coordinates and fixing alleles of the PsychENCODE genotypes to the hg38 reference genome, we performed strand-flipping using the snpflip software package [40]. We removed SNPs with minor allele frequency <0.05 or Hardy-Weinberg equilibrium *p* < 1 *×* 10^−6^ (resulting in more than 2.6M SNPs per sample) and removed individuals with >3 SD genotype heterozygosity rate (F coefficient, calculated using SNPs in 200 kb windows with LD *r*^2^ > 0.25). An additional strand-flipping step was performed using snpflip to match alleles between the sample genotypes and summary statistic SNPs. Finally, six genotype principal components (PCs) were calculated for each sample using the –pca command in PLINK2 [41].

We used the LDpred2 function within the bigsnpr software package to calculate PRSs for each individual [42]. After filtering for SNPs present in the HapMap3 dataset, we calculated LD scores for 110,258 matching SNPs between the PsychENCODE genotypes and summary statistics using the snp cor function and centimorgan map units from the 1000 Genomes project. We then regressed the LD scores against the log-scaled summary beta values using the snp ldsc function and calculated a baseline heritability estimate based on the output (*h*^*2*^ *_est*, here equal to 0.1270). Using these modified summary statistics, we calculated PRSs for each individual using the infinitesimal and grid models of LDpred2. For the grid model, we assessed combinations of (a) 17 input SNP *p*-value filters ranging from 1.0×10^-4^ to 1.0, (b) three potential values of *h*^*2*^ (0.7, 1.0, and 1.4×*h*^*2*^ *_est*), and (c) sparse or non-sparse grid models, for a total of 102 PRS outputs. (We note that the LDpred2-auto model did not converge for our datasets and thus was not used in our analysis.) To assess the performance of each PRS model, we generated logistic regression models for PRSs towards the binary schizophrenia diagnosis of each individual, with sex, age, and six genotype PCs included as covariates. We then calculated Nagelkerke *pseudo-R*^*2*^ values for each regression model, and compared them with those for a null logistic regression model consisting solely of the sex, age, and genotype PC covariates.

Overall, we found that both the infinitesimal model (*pseudo-R*^*2*^=0.2008) and best-performing grid model (*pseudo-R*^*2*^*=*0.2229; at *p*=0.32, *h*^*2*^=0.7×*h*^*2*^ *_est*, and sparse grid model) performed substantially better than the null model (*pseudo-R*^*2*^=0.0809). We selected PRS outputs derived from the best-performing grid model for comparison with encrypted PRS calculations. The correlations and AUROC calculations performed in this analysis were completed in Python using the sklearn library.

In contrast to the LDpred2 model, our PRS calculation approach used for the homomorphic encryption protocol performs SNP filtering as a preprocessing measure. In this way, we ensure that the input to our model consists of already filtered SNPs. Additionally, our model does not explicitly manage LD in the same detailed manner. Instead, it operates similarly to a straightforward PRS model, which assumes that the selected SNPs are representative of all SNPs that contribute to a phenotype and have been pre-processed to mitigate LD concerns.

## 4 Results

### 4.1 Implementation Results

We evaluated the performance of our FHE-based PRS calculation method using both synthetic datasets and a real dataset for schizophrenia risk. We first applied our method to a synthetic dataset created using Hapgen2 [35], which contained 2,000 individuals and a range of SNPs for three artificial phenotypes (referred to as phenotypes 0, 1, and 2). We then applied our method to a dataset of over 1,100 individuals from the PsychENCODE Consortium [38], evaluated against a 110K–SNP PRS model for schizophrenia [28]. We compared the performance of our encrypted PRS with the non-encrypted PRS in terms of score correlation (Pearson r) and mean squared error (MSE) in phenotypic variance explained by the scores.

#### 4.1.1 Overview of PRS Derivations

We used two complementary approaches to assess the performance of our homomorphic encryption-based PRS calculations. We first generated synthetic datasets using Hapgen2 [35], a simulation method that produces genotypes with realistic LD patterns, to further evaluate our approach under various conditions. We simulated datasets with varying numbers of SNPs (10k to 130k) and sample sizes (200 to 2,000 individuals) and calculated PRSs for three artificial phenotypes with different levels of heritability using ridge regression. By assessing our FHE method on both real and synthetic data, we demonstrated its versatility and potential for secure and private PRS calculations in diverse scenarios. We also applied our method to a real-world dataset for schizophrenia, utilizing PRSs derived from a recent GWAS of the PGC of more than 320,000 individuals [28]. Specifically, we calculated PRS for 1,146 individuals (493 cases and 653 controls) from the PsychENCODE Consortium [38], using a standardized data processing pipeline based on best practice guidelines [39], followed by LDpred2 for PRS calculation [42]. The resulting PRS model contained 110,258 SNPs and achieved a pseudo-R^2^ value of 0.2229 in predicting schizophrenia diagnosis.

For both approaches, we found that our FHE procedure introduced only a trivial amount of error compared to the inherent uncertainty of the PRS value as determined from non-encrypted approaches, despite the framework’s computational and memory limitations with increasing data size. Specifically, the variability of the phenotype explained by PRS (*R*^2^) from our homomorphic encryption framework closely matched that of the underlying PRS models for both synthetic and real data, indicating that the explained variance in the phenotype within the population remained consistent after encryption. For example, we observed a small MSE (<1.5 *×* 10^−8^) when applying FHE with varying *R*^2^ values to the synthetic datasets (Table 3). After ridge regression analysis on the encrypted data, the *R*^2^ values obtained for the three artificial phenotypes were 0.189, 0.466, and 0.822, respectively (compared with the preset values of 0.3, 0.6, and 0.9). Similarly, using the 110K SNP dataset for schizophrenia PRS, our encrypted PRS values had an *R*^2^ value of 0.2232 (with PRS, sex, age, and genotype PCs as covariates), compared with 0.2226 for the non encrypted model (Supplementary Table 5). This corresponded to an MSE of 2.27 × 10^−6^ between the non encrypted PRS and the encrypted PRS (for ring dimension 2^13^). This minimal MSE demonstrated that the encrypted PRS data yielded comparable *R*^2^ values to those obtained without encryption, suggesting that our FHE method introduces only a small error that would not lead to changes in the derived PRS. Figure 3A demonstrates that the LDpred2 and encrypted PRS methods both achieved an AUROC of 0.61. Additionally, Figure 3B shows a Pearson correlation of *r* > 0.999 between the encrypted and non-encrypted schizophrenia PRS. In terms of computational cost, Figure 3C shows that the majority of time is spent on input encryption and encrypted computation, while model encryption and result decryption require comparatively little time.

**Table 3:** Performance of encrypted PRS models with ring dimension 2^13^ across three phenotypes with varying heritability (h^2^ = 0.3, 0.6, 0.9). The R^2^ of the encrypted model and MSE relative to the corresponding plaintext model are shown.

| Phenotype | Ring<br>Dimension | $R^2$ of plaintext<br>model | $R^2$ of encrypted<br>model | MSE between<br>plaintext and<br>encrypted model |
| --- | --- | --- | --- | --- |
| 0 | $2^{13}$ | 0.18892353 | 0.18892715 | 1.397e-08 |
| 1 | $2^{13}$ | 0.46645884 | 0.4664542 | 8.085e-09 |
| 2 | $2^{13}$ | 0.8226163 | 0.8226161 | 1.168e-08 |

**Table 4:** Accuracy comparison of PRS generated from plaintext ridge regression models and encrypted models. Data were generated using Hapgen2, including three phenotypes corresponding to different levels of SNP heritability (h2=0.3, 0.6, 0.9), which we refer to as phenotype 0, 1 and 2. The R^2^ between simulated phenotypes and PRS predicted by the plaintext and encrypted models are shown. MSE values are shown to compare the predictions with and without encryption. Statistics shown derive from a model with 10,000 SNPs and 2,000 individuals.

| Phenotype | Ring<br>Dimention | $R^2$ of plaintext<br>model | $R^2$ of encrypted<br>model | MSE between plaintext<br>and encrypted model |
| --- | --- | --- | --- | --- |
| 0 | $2^{12}$ | 0.18892353 | 0.1889256 | 1.59e-09 |
| | $2^{13}$ | | 0.18892715 | 1.397E-08 |
| | $2^{14}$ | | 0.18892446 | 7.778E-10 |
| | $2^{15}$ | | 0.18892441 | 0 |
| | $2^{16}$ | | 0.18892443 | 0 |
| 1 | $2^{12}$ | 0.46645884 | 0.4664603 | 7.088e-09 |
| | $2^{13}$ | | 0.4664542 | 8.085E-09 |
| | $2^{14}$ | | 0.46646012 | 1.920E-10 |
| | $2^{15}$ | | 0.46645956 | 0 |
| | $2^{16}$ | | 0.46645956 | 0 |
| 2 | $2^{12}$ | 0.8226163 | 0.8226161 | 1.878e-09 |
| | $2^{13}$ | | 0.82261809 | 1.168E-08 |
| | $2^{14}$ | | 0.82261557 | 1.426E-09 |
| | $2^{15}$ | | 0.82261601 | 0 |
| | $2^{16}$ | | 0.822616 | 0 |

**Figure 3:**
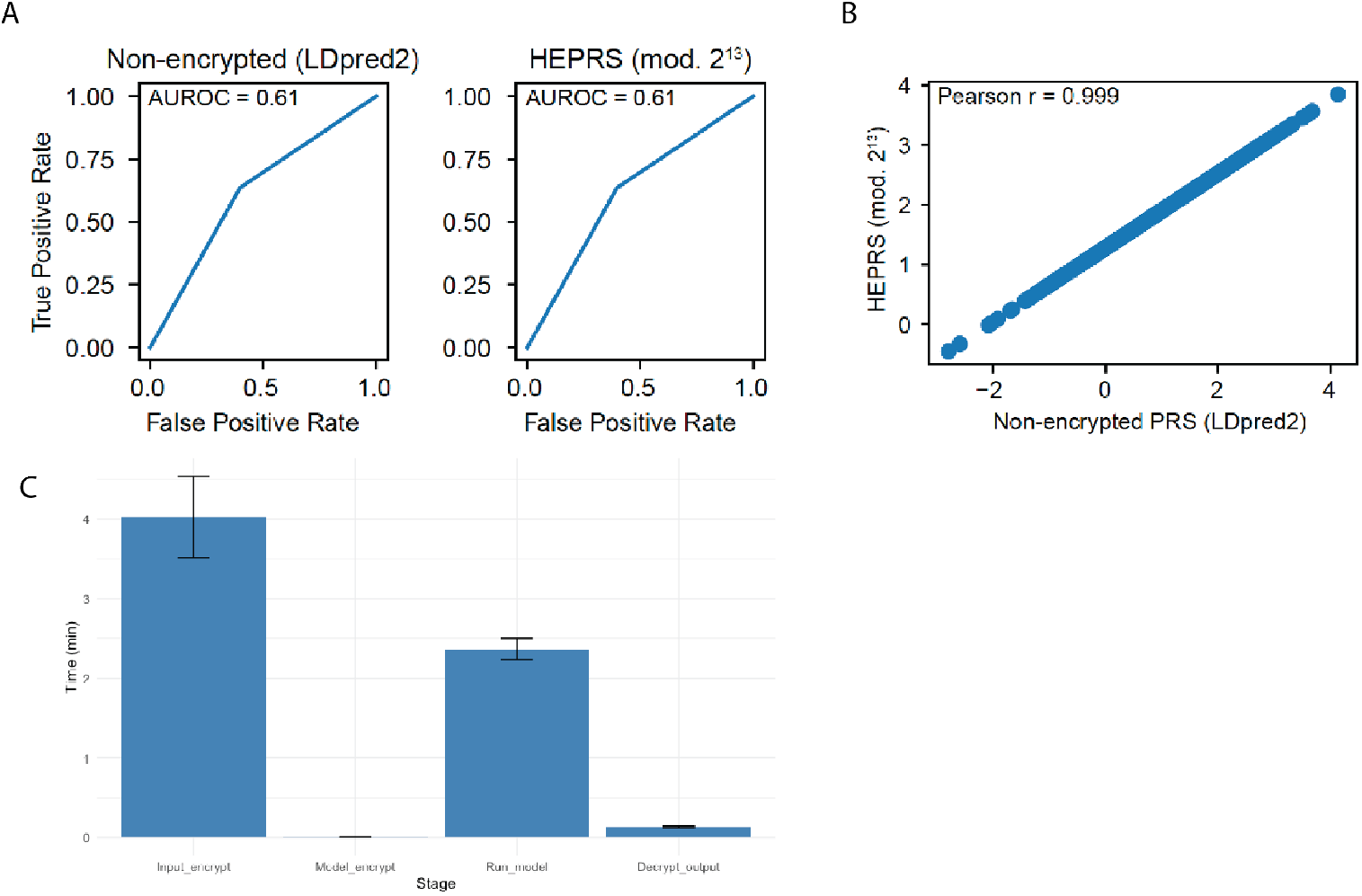
(A) Receiver operating characteristic curves for prediction of schizophrenia phenotypes based on non-encrypted schizophrenia PRSs (calculated with LDpred2) and PRSs calculated from HEPRS using the ring dimension parameter 2^13^. PRS accuracy was assessed by considering individuals with > 0.5 percentile scores for each calculation to have schizophrenia, and comparing with the actual phenotypes. AUROC values were equal across all methods. (B) Pearson correlations between encrypted schizophrenia PRSs with the ring dimension parameter 2^13^ and non-encrypted PRSs calculated with LDpred2 each show a high correlation (r > 0.999). (C) Break-down of computation time into different steps of the homomorphic encryption process using the ring dimension parameter 2^13^. The time was calculated by assessing PRSs of 1,146 individuals using the 110K SNP schizophrenia model on a 6234 CPU.

### 4.2 Effects of Input Data Size on Runtime and Memory for Encrypted PRS

While several parameters may affect runtime and memory usage, our method was able to quickly and accurately calculate PRSs using encrypted genotypes and PRS models, even when implemented on a personal laptop. For instance, using a MacBook Pro (M1, 2020), our method required about 10 seconds and 3 GB of memory to calculate PRSs for one individual sample using the 110K schizophrenia PRS model (ring dimension 2^13^). Calculating PRSs for a larger cohort of 1,146 samples on a single CPU (Intel 6234) required 6 minutes and about 65 GB of memory when using the 110K SNP model with the ring dimension 2^13^ (Supplementary Table 5). The run time for 2,000 samples in the 10K SNP synthetic dataset took about 2 minute for any of the three synthetic phenotypes..

Supplementary Table 5 demonstrates that the protocol utilizing homomorphic encryption for dot product calculation was roughly three orders of magnitude slower than plaintext calculations (on a single CPU node Intel 6234). When we implemented the dot product calculation in Golang, it took 0.303 seconds; however, a more common way to calculate the PRS is with LDpred2 in R, which took only 0.073 seconds. When comparing the dot product calculation method within Golang, the homomorphic encryption calculation (248.3 seconds) took around 800 times longer. Compared to standard methods of calculating the PRS, the encrypted calculation took 3,400 times longer. This contrast highlights the computational burden introduced by encryption protocols and the possible time added by the Golang language.

Using a synthetic dataset allowed us to vary both the input sample size and the number of SNPs of the underlying PRS model to determine their effects on the performance of our method.

We first assessed the influence of the number of SNPs on the time taken to calculate PRSs using FHE for 50 samples (Fig. 4). We observed a direct correlation between the increase in SNP numbers, from 10,000 to 130,000, and the computational time required, from 5 seconds to 36 seconds, demonstrating an approximately linear relationship under the conditions tested (Fig. 4 A). Figure 4B expands on this by showing that memory requirements also rise in a similar linear fashion (from 0.8 GB to 9 GB) with the increase in SNP count, which is a critical consideration for the practical application of FHE in clinical settings.

**Figure 4:**
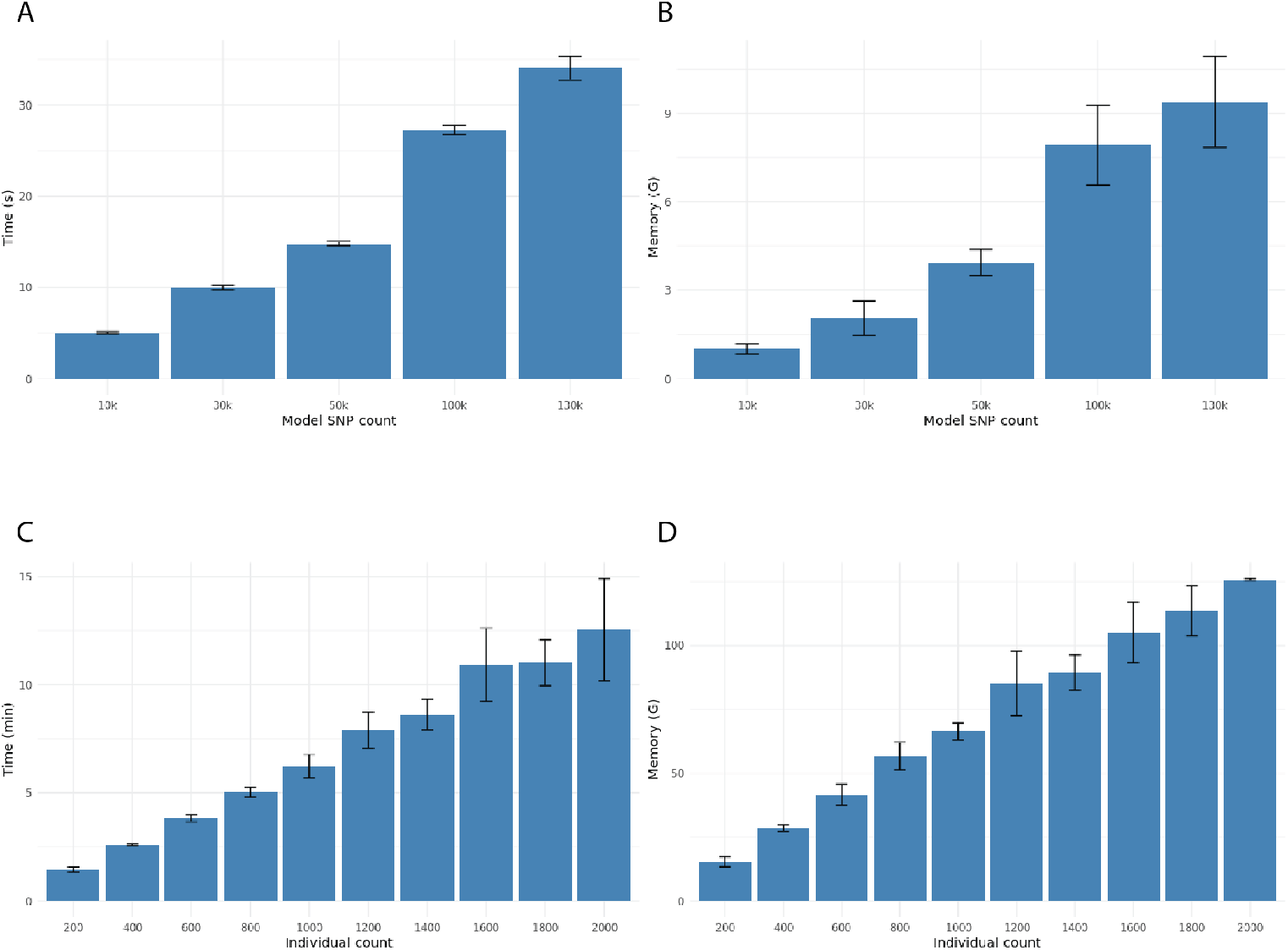
Homomorphic encryption computation time and memory usage scale approximately linearly with the number of SNPs and individuals. (A) Computation time and (B) memory usage as a function of the number of SNPs, generated using synthetic datasets containing a range of SNP sizes created by Hapgen2. The runs were performed using a ring dimension 2^13^, a sample size of 50 individuals, and phenotype 0 (as specified in Table 1 and Table 4). (C) Computation time and (D) memory usage as a function of the number of individuals, using the synthetic dataset with 130,000 SNPs and varying only the number of individuals for each run. A ring dimension of 2^13^ and phenotype 0 were used for these experiments. Error bars indicate the standard deviation from 10 independent trials tested on a random choice from a 6234 CPU.

Next, we varied the number of individuals for which we performed the encrypted PRS calculations, using a constant genome size of 130K SNPs and predicting for artificial phenotype 0. Similar to the number of input SNPs, we found that the number of individuals included in a batch had a linear relationship with the time to completion (Fig. 4C), ranging from about 1.5 minutes for 200 individuals to 12.5 minutes for 2,000 individuals. The required memory also positively correlated with the batch size, from 17 GB for 200 individuals to 125 GB for 2,000 individuals (Fig. 4D). These data point to a predictable increase in analysis time with larger cohorts, reinforcing the scalability of our FHE protocol for PRS calculation in real-world scenarios with substantial genomic datasets.

## 5 Discussion

In this work, we implemented an FHE-based method for accurate and efficient PRS calculations without compromising patient privacy. We show that the marginal error introduced by the encryption scheme does not affect the overall error of the PRS model. We also demonstrate that different parameters affect the accuracy of results and the trade-off between accuracy and time within the FHE framework. Although FHE substantially increases the computational burden for PRS calculation compared to plaintext models, roughly three orders of magnitude slower, the cost remains within a reasonable range for practical applications. These findings suggest that our FHE-based method is viable for securely computing PRSs while maintaining a balance between privacy, accuracy, and computational efficiency.

The bulk of our PRS calculation is performed within our function `HE_inner_product()`. While this operation is a simple calculation, the nature of large genomic data requires some reworking of encrypted data storage and the computation sequence to arrive at an accurate phenotype prediction without excessive time or memory consumption. In the Methods section, we provide an intuitive explanation of how we utilize RLWE to compute the dot product.

As explained in the Methods, the modeler must share the encrypted model with the evaluator; if the evaluator were to share this encrypted model with the client, the client could decrypt the model without the modeler’s consent, thus violating privacy requirements. The assumption that the evaluator and client never collude is reasonable if we consider the evaluator to be a healthcare entity (e.g., a hospital or healthcare insurance institution). These organizations might be associated with a research group that owns the model. We assume the evaluator would want to maintain a good relationship with the modelers and not erode trust. However, the trust between the modeler and evaluator poses a vulnerability in the framework.

Approximately 4–7% of an average healthcare system’s IT budget is spent on cybersecurity. This means that from 2020 to 2025, healthcare organizations will allocate $125B to cybersecurity [43]. While hospitals use a large portion of this budget to defend the Internet of Things, some of this capital is spent protecting healthcare data, which increasingly includes patient genomic data. Our protocol could significantly reduce vulnerabilities if patient genomes were stored in encrypted formats and only the decryption keys were maintained under high security. Such an encryption framework would reduce vulnerabilities because only encrypted genomic data would be transferred, rather than raw genomic data. However, privacy could still be breached if the evaluator and client colluded to reveal the underlying PRS weights.

As our pipeline assumes a single decryption authority (the client), we use one public/secret key pair shared by all parties for encryption and evaluation. This is optimal when only the genotype owner needs access to the plaintext result and the PRS model owner (researcher) trusts the evaluator. Specifically, our protocol involves three logical roles (client, modeler, evaluator), all operating under a single CKKS public key generated by the client (Fig. 1). This “single-key, many-party” layout is a natural match for the encrypted inference use case we target: the client (e.g., a clinician) requests phenotype predictions on a patient’s genotype without revealing raw variants to the cloud evaluator; the evaluator and model provider do not receive access to genotypic information; and the evaluator cannot decrypt the model or genotype. Thus, this method relies on the assumption that the evaluator and client will not collude to reveal the model parameters and assumes trust between the evaluator and modeler. However, many genomic workflows involve multiple mutually distrustful custodians, such as a consortium of hospitals storing data for a subset of patients, or two companies wishing to jointly train a PRS without exposing proprietary cohorts. In such cases, a multiparty (collective-key) CKKS scheme would remove the need to trust any single organization with the full secret key. In this context, the secret key is additionally shared across the parties; no single party can decrypt, but a quorum can run a threshold-decryption (or key-switch) protocol. This mitigates the “evaluator and modeler trust” assumption at the cost of increased interaction and computation. True multiparty CKKS (collective-key or MPC-FHE) requires interactive key generation and online key-switch refreshes, introducing >3*x* communication overhead. This would add runtime and complexity, which is unnecessary in our current context where only the client requires decryption capability. However, a multiparty-FHE PRS pipeline is a possible next step and could be valuable in different contexts where a federated learning process is desired, such as when there are multiple modelers or when the model provider and evaluator do not trust each other.

In this work, we demonstrated the feasibility and practicality of generating PRS model predictions without revealing a genome. We showed how memory requirements increase with higher accuracy specifications, and how runtime scales roughly linearly with the number of SNPs included in the model. Lastly, we showed that the error introduced by homomorphic encryption is negligible compared to the inherent error in the model itself.

## 6 Conclusion

Considering recent advances in FHE, we view PRSs as an exciting new avenue for applying this technology. The strategic use of genomic information has the potential to significantly contribute to personalized healthcare to enhance health outcomes and elevate care standards. The necessity for patients to undergo genomic sequencing is paramount for realizing this technology’s benefits. Concurrently, the escalation of genetic data mandates stringent security and privacy measures. The challenges of applying analytical models to decipher relevant information from the large-scale data required for genome encoding are non-trivial. Our research offers a solution that addresses these multifaceted issues. While our focus is on PRSs, the principles of FHE can potentially be applied to a spectrum of genomic computations. The generalizability of FHE to other areas in clinical genomics, such as variant annotation, genotype imputation, and more complex predictive modeling, warrants further investigation. This work takes an initial step toward exploring such applications.

## List of abbreviations

FHE: Fully Homomorphic Encryption
PRS: Polygenic Risk Score
CKKS: Cheon-Kim-Kim-Song
SNP: Single-Nucleotide Polymorphism
GWAS: Genome-Wide Association Studies
PGC: Psychiatric Genomics Consortium
MSE: Mean Squared Error
RLWE: Ring Learning With Errors
LDpred2: LDpred version 2
PC: Principal Component

## Declarations

### Ethics approval and consent to participate

Not applicable.

### Consent for publication

Not applicable.

### Availability of data and materials

The datasets generated and/or analyzed during the current study are available from the corresponding author upon reasonable request. PsychENCODE genotypes are available for download by approved researchers at http://resource.psychencode.org/, and GWAS summary statistics for schizophrenia are available from the Psychiatric Genomic Consortium portal at https://pgc.unc.edu/for-researchers/download-results. Code is available as a GO package at https://github.com/gersteinlab/HEPRS, along with a small example dataset that includes synthetic genotypes and phenotypes generated using Hapgen2 for tutorial purposes.

### Authors’ contributions

## Authors’ information

## Supplementary Materials

### 6.1 Ring Dimension Affects Accuracy and Runtime for PRS Calculation

We showed above that increasing the ring dimension N parameter of the CKKS encryption scheme will decrease the errors introduced in the encrypted calculations. However, increasing the ring dimension will also increase the model runtime.

From the results obtained with the 110K–SNP schizophrenia PRS model, we observed that the time required for the encrypted model to generate PRS predictions increased with the ring dimension (Fig. 5A). Evaluating PRSs for 1,146 individuals using the 110K–SNP model in plaintext (infinitesimal mode of LDpred2 for only the evaluation part) typically takes 0.07 seconds on a single CPU (Intel 6234) (Table 5), while for FHE, even using the smallest ring dimension 2^13^ takes about 6 minutes (Fig. 5A), and the time further increases to 20 minutes for the ring dimension 2^16^. When breaking down the time cost into different stages of the encryption calculation, we find that most of the time is spent after the encryption step and during the model operation phase, corresponding to the actual computation of the encrypted PRS (Fig. 5B).

**Table 5:** Comparison of PRSs generated from the encrypted model and LDpred2 model. The coefficient of determination (R2) for LDpred2 and encrypted models were generated from fitting logistic regression models towards the binary schizophrenia diagnosis of each individual, with PRS, sex, age, and six genotype PCs included as covariates. The Nagelkerke pseudo-R2 values for each regression model are reported. The R2 between encrypted and LDpred2 models were calculated from a linear regression between encrypted and unencrypted PRSs. The MSE was also calculated between encrypted and unencrypted predictions. The time for running our encrypted model is listed for each specified ring dimension. The input consists of 1,146 individuals with 110,258 SNPs. Time reported for homomorphic encryption calculation is the mean ± standard deviation record from 10 independent runs. Times for HE, LDpred2, and plaintext Golang (with and without reading and writing) calculations were all performed using a single CPU chosen randomly from a 6234, 6240 or 6346 CPU.

| Ring Di-<br>mention | R<sup>2</sup> of<br>LDPred2<br>model | R<sup>2</sup> of<br>encrypted<br>model | R2 between<br>encrypted<br>and<br>LDPred2<br>model<br>predictions | MSE<br>between<br>LDPred2<br>and<br>encrypted<br>model | Time for<br>LDPred2<br>Infinitesi-<br>mal model<br>calculation<br>(s) | Time for<br>HE<br>calculation<br>(s) [with<br>reading and<br>writing] | Time for<br>HE<br>calculation<br>(s) [without<br>reading and<br>writing] | Time for<br>plaintext<br>calculation<br>in Golang<br>(s) [with<br>reading and<br>writing] | Time for<br>plaintext<br>calculation<br>in Golang<br>(s) [without<br>reading and<br>writing] |
| --- | --- | --- | --- | --- | --- | --- | --- | --- | --- |
| 2<sup>12</sup> |  | 0.2233236 | 0.9999999 | 1.158E-07 |  | 303.53 ±<br>0.5849 | 182.7721 |  |  |
| 2<sup>13</sup> |  | 0.2232212 | 0.9999895 | 2.268E-06 |  | 391.98 ±<br>34.85 | 248.3242 |  |  |
| 2<sup>14</sup> |  | 0.2233313 | 1.0000000 | 1.430E-06 |  | 498.30 ±<br>36.61 | 319.1728 |  |  |
| 2<sup>15</sup> |  | 0.2233307 | 1.0000000 | 1.425E-06 |  | 669.48 ±<br>30.07 | 469.8914 |  |  |
| 2<sup>16</sup> | 0.2226318 | 0.2233310 | 1.0000000 | 1.425E-06 | 0.073 | 1233.64 ±<br>32.09 | 1020.181 | 6.224 | 0.303 |

**Figure 5:**
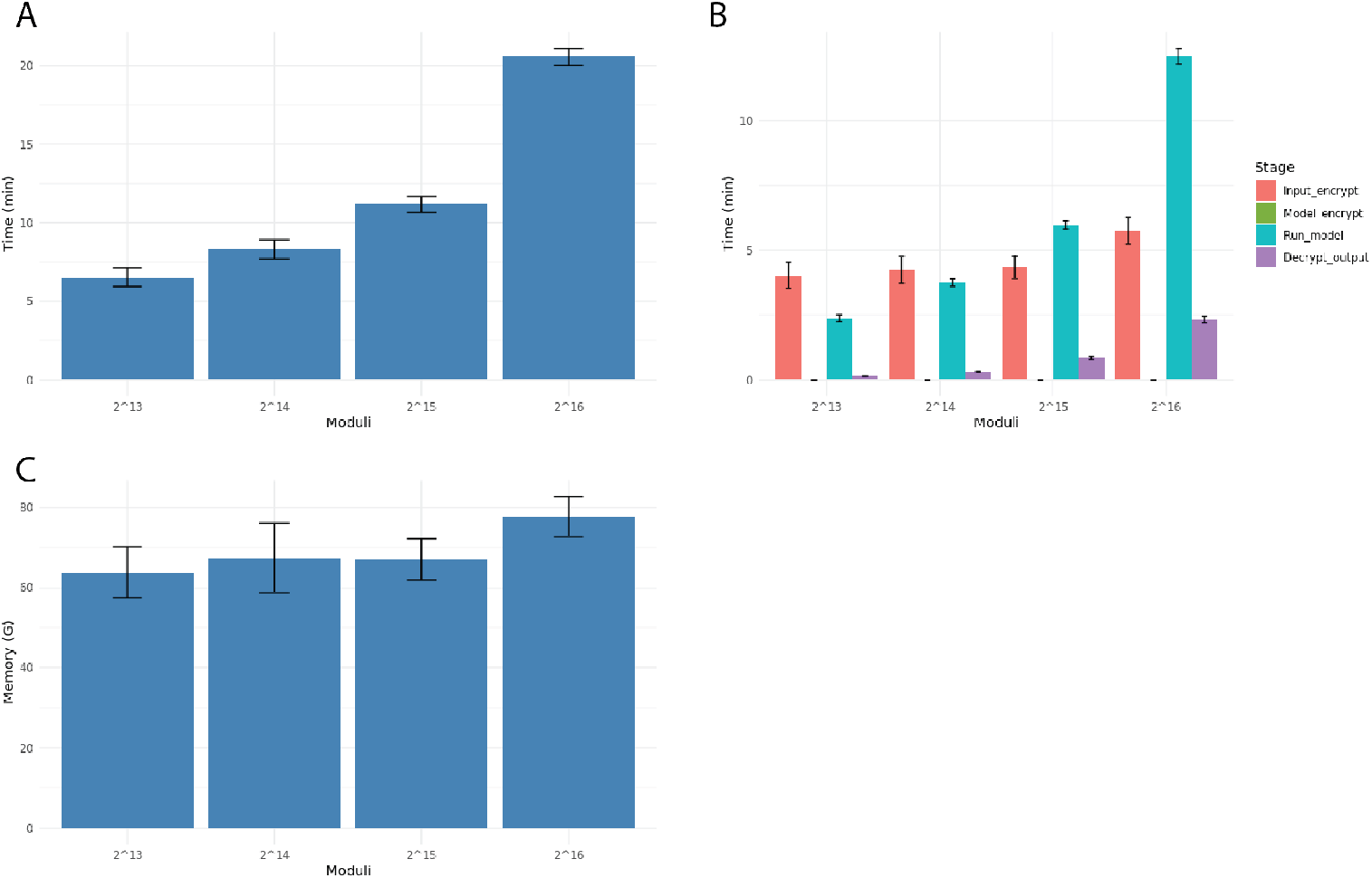
The effect of ring dimension on computation time and memory usage in homomorphic encryption for a schizophrenia PRS model generated from real data. The model contains 1,146 individuals and 110,258 SNPs. The range of tested ring dimension is from 2^13^ to 2^16^. (A) Total computation time as a function of ring dimension. (B) Breakdown of computation time into different steps of the homomorphic encryption process. (C) Memory usage as a function of ring dimension. Error bars indicate the standard deviation calculated from 10 independent trials tested on a 6234, 6240, or 6346 CPU.

Fig. 5C presents the memory allocation for running the program across different ring dimensions. Contrary to what we expected, the memory usage remains largely invariant through repeated experiments, ranging from 60 GB to 80 GB, which only introduces a moderate increase in memory usage compared to the plaintext model, which is 50 GB. Thus, our FHE method for PRS calculation allows end users to account for the trade-off between protocol time and error mitigation according to their needs by customizing the ring dimension parameter.

For both synthetic and real datasets, we observe a decrease in MSE as we increase the ring dimension N of the encryption scheme from 2^13^ to 2^16^, which suggests a reduction in computational error with higher ring dimension. For example, the MSE reduces to 1.43 × 10^−6^ for the real dataset with a ring dimension of 2^16^ (Table 5). This suggests that a higher ring dimension can mitigate computational errors introduced by the encryption. Furthermore, we show in Figure 6A that all calculation methods maintain a nearly constant AUROC. In Figure 6B, we show a Pearson correlation with r >0.999 between the encrypted and non-encrypted schizophrenia PRS regardless of the ring dimension.

**Figure 6:**
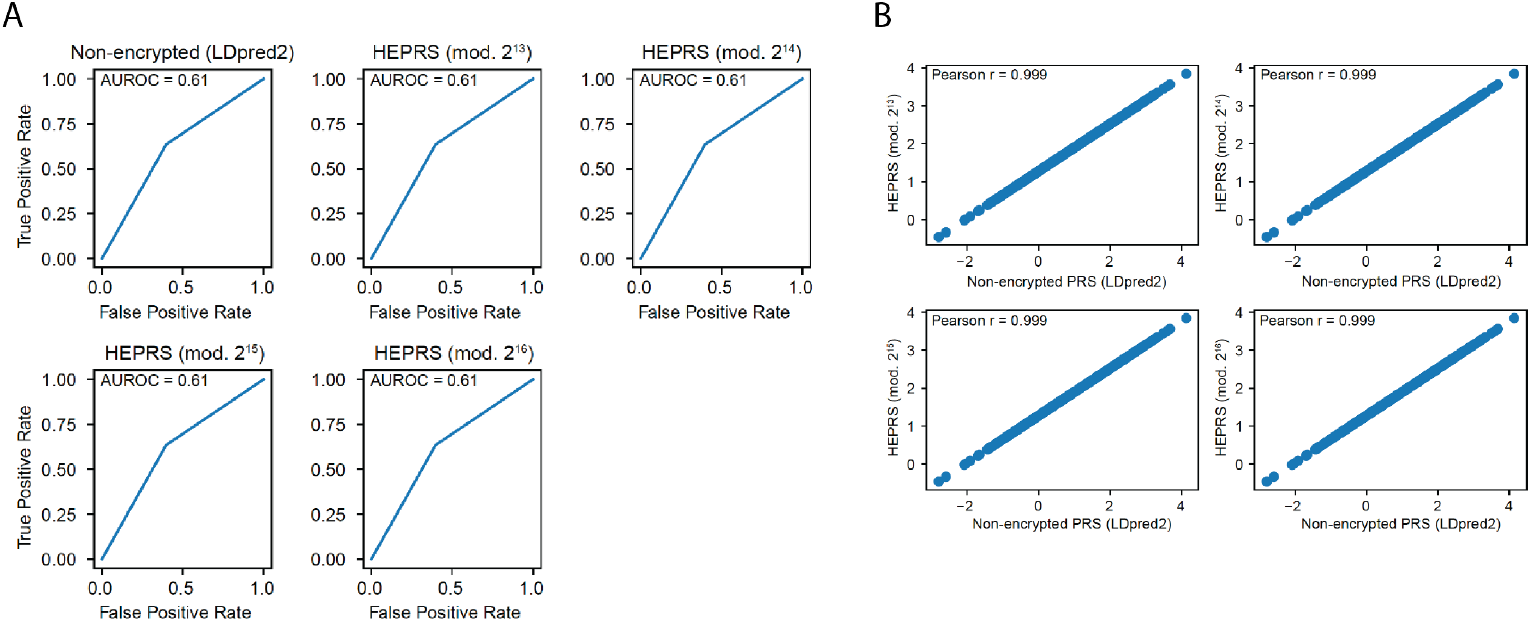
(A) Receiver operating characteristic curves for prediction of schizophrenia phenotypes based on non-encrypted schizophrenia PRSs (calculated with LDpred2) and PRSs calculated from HEPRS using four different ring dimension parameters. PRS accuracy was assessed by considering individuals with > 0.5 percentile scores for each calculation to have schizophrenia, and comparing with the actual phenotypes. AUROC) values were equal across all methods. (B) Pearson correlations between encrypted schizophrenia PRSs with four different dimension parameters and non-encrypted PRSs calculated with LDpred2 each show a high correlation (r > 0.999).

